# Correlation of mRNA delivery timing and protein expression in lipid-based transfection

**DOI:** 10.1101/607986

**Authors:** A. Reiser, D. Woschée, N. Mehrotra, R. Krzysztoń, H. H. Strey, J. O. Rädler

## Abstract

Non-viral gene delivery is constrained by the dwell time that most synthetic nucleic acid nanocarriers spend inside endosomal compartments. In order to overcome this endosomal-release bottleneck, methods are required that measure nanocarrier uptake kinetics and transfection efficiency simultaneously. Here, we employ live-cell imaging on single-cell arrays (LISCA) to study the delivery-time distribution of lipid-based mRNA complexes under varied serum conditions. By fitting a translation-maturation model to hundreds of individual eGFP reporter fluorescence time courses, the protein expression onset times and the expression rates after transfection are determined. Using this approach, we find that delivery timing and protein expression rates are not intrinsically correlated at the single-cell level, even though population-averaged values of both parameters conjointly change as a function of increasing external serum protein fraction. Lipofectamine mediated delivery showed decreased transfection efficiency and longer delivery times with increasing serum protein concentration. This is in contrast to ionizable lipid nanoparticles (LNPs) mediated transfer, which showed increased efficiency and faster uptake in the presence of serum. In conclusion, the interdependences of single-cell expression rates and onset timing provide additional clues on uptake and release mechanisms, which are useful for improving nucleic acid delivery.

## Introduction

Gene based therapies advance the delivery of exogenous nucleic acids with the intent to modulate the expression of disease related genes. However, clinical applications are still limited due to the difficulties inherent in delivery of nucleic acids *in vivo* (1, 2). Over the last years, intense research is dedicated to develop non-viral vectors that are capable to safely and efficiently package, transport and systemically deliver nucleic acid to cells (3-5). In particular for therapeutic purposes, there is need to deliver DNA, mRNA, small interfering RNA (siRNA) or microRNA (miRNA). The molecular and cellular processes of nucleic acid nanoparticles share common structural features and entry mechanisms. Some of the major challenges associated with current cationic lipid vectors are the interaction with blood serum components, impaired intracellular uptake, limited release into the cytosol and immunological response (6, 7). Each of these hurdles are independently affected by variations in the lipid formulations and transfection conditions. Lipid-encapsulated RNA is taken up by various, endosomal or phagocytotic, uptake mechanisms and subsequent intracellular pathways. It appears however, that nucleic acids nanocarriers are largely trapped inside the endosomes or lysosomes and that delivery by lipid nanoparticles is limited by endocytic recycling (8). In time-resolved fluorescence microscopy studies, the uptake and fate of delivery particles has been visualized at the single-cell level (9-13). Recently, it was shown that siRNA release occurred during a narrow ‘window of opportunity’ showing that timing and efficiency are linked (12). However, the uptake pathways and factors that control endosomal release of lipid-based nanocarriers are not fully resolved. In particular, the interdependence of timing and efficiency of gene delivery and the role of extracellular factors on the kinetics of cellular uptake and endosomal release are poorly understood aspects in the delivery process.

Recently, we established quantitative live-cell imaging on single-cell arrays (LISCA) for the time-resolved study of reporter gene expression at the single-cell level (14-16). In case of mRNA-mediated eGFP expression, the single-cell fluorescence was found to follow a first-order translation model, which renders the kinetics of mRNA expression with high predictive precision (15, 16). Among others, the single-cell analysis infers the individual expression onset times after mRNA transfection. These findings open the door to establish a routine assay to assess the delivery timing in terms of a distribution of individual time-to-expression after transfection (17). Furthermore, we automated single-cell fluorescence readout using micro-structured substrates with arrays of single-cell adhesion sites combined with a tailored perfusion system enabling fluid exchange for mRNA transfection during the time-lapse measurement (18). The platform provides access to the distribution of expression onsets and expression rates after transfection. In this context, a systematic study of nanocarrier uptake kinetics, the expression onset times, and how it relates to protein expression efficiency and external transfection conditions has not yet been carried out at the single-cell level.

Here we study the time-to-expression after mRNA delivery as well as the expression rate using high-throughput single-cell analysis of eGFP reporter fluorescence. We employ micropatterned single-cell arrays and automated live-cell time-lapse microscopy to collect individual fluorescence time courses. Fitting the time courses to a translation-maturation model, we obtain best estimates for the onset time and expression rate distributions. The single-cell data reveal large cell-to-cell variability and the insight that the delivery time and expression rate do not correlate at the single-cell level. Nevertheless, both parameters change in a concerted manner as a function of external parameters such as the amount of serum. In mRNA transfection experiments using Lipofectamine lipoplexes the efficiency decreases, and delivery timing increases as a function of the fraction of serum protein. In contrast, for lipid nanoparticles (LNPs) containing the ionizable cationic lipid DLin-MC3-DMA efficiency improves and timing shortens in the course of increasing serum.

## Results

### Highly parallel single-cell protein expression readout by scanning time-lapse microscopy

In order to record single-cell protein expression time courses in a highly parallel manner we employed an LISCA approach as published earlier (18). HuH7, liver carcinoma, cells were seeded onto a microstructured cell culture channel slide (see Figure 1A). In order to capture the very early events of protein expression the single-cell array was connected to a tailored perfusion system and scanning time-lapse data acquisition was started prior to transfection. The perfusion system allowed for fluid exchange and enabled the addition of transfection agents on the stage during the time-lapse measurement. mRNA complexes were added one hour after the beginning of the measurement and incubated for one hour. After the incubation period the transfection solution was exchanged for cell growth medium (indicated by the arrows and the gray bar in Figure 1B. Note that without the perfusion system, it is not possible to observe changes in the fluorescence intensity during and shortly after the time period of the transfection due to sample handling off the microscope. Secondly, with the use of a perfusion system, we prevent particle adsorption events after the incubation time by flushing out unbound nanocarriers. This is illustrated in Supplementary Figure 1A using fluorescently labeled lipoplexes before and after flushing. To investigate whether nanocarrier are taken up, we prepared lipoplexes containing mRNA with a Cy5 labeled fraction to visualize the lipoplexes during the time-lapse measurement. The fluorescence kinetics of the Cy5 labeled mRNAs shows the adsorption onto the cell surfaces (see Supplementary Figure 1B). As a reference, we observed on areas with no cells that the fluorescence time courses of nanocarriers increases during the incubation period and abruptly decreases when the chamber is flushed with fresh carrier-free medium indicating that some partially adherent nanocarriers are rinsed away. Interestingly, in contrast, the fluorescence of nanocarriers adsorbed to cells not only does not decrease but increases within the next 1 hour after flushing. We interpret this time course such that firstly, nanocarriers are strongly adsorb and are not rinsed away and secondly, the fluorescence of the Cy5 labeled mRNAs is likely to increase during the time of uptake due to unpacking of the lipid-based carriers and thus dequenching the Cy5 fluorescence.

**Figure 1:**
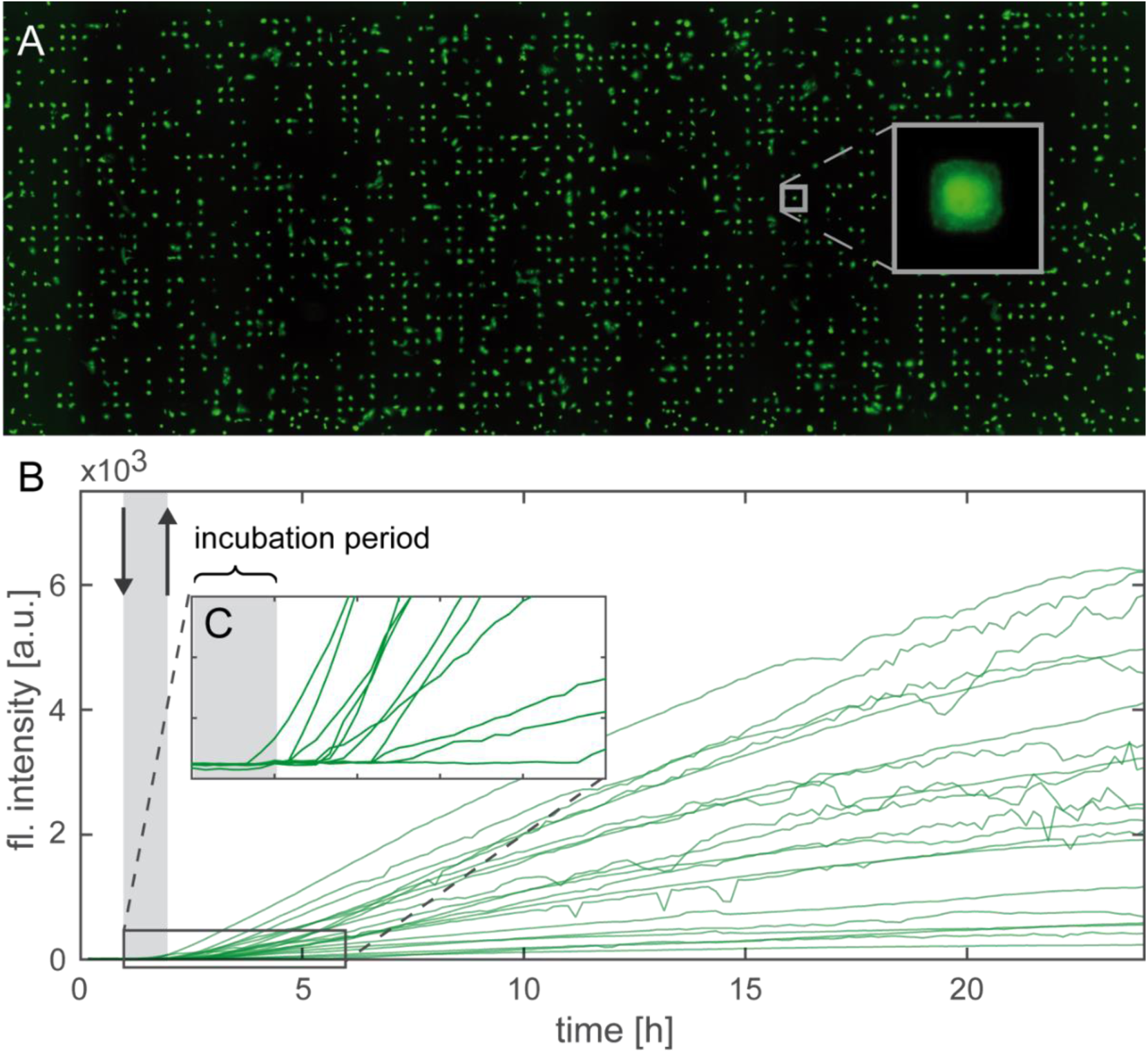
Highly parallel single-cell measurement of eGFP protein expression after mRNA transfection show large cell-to-cell variability. Microstructered single cell arrays are comprised of a protein square pattern on which the cells preferentially adhere. This confinement results in standardized conditions for each cell (A). The cells are transfected with mRNA by adding mRNA nanocarriers using a perfusion system and incubating the complexes for one hour before the fluid is exchanged with cell growth medium (the two perfusion steps and the incubation is indicated by the arrows and the gray bar). Subsequently, we record the fluorescent protein expression time courses. Representative single-cell time courses illustrate the variation for protein expression levels (B) and eGFP expression onset times (C).

In the first set of experiments Lipofectamine2000 was used as transfection agent and mRNA encoding for eGFP as reporter (see Materials and Methods). The scanning time-lapse measurement sequentially yields fluorescence image stacks from typically 70-100 positions. The movies are further processed to retrieve single-cell protein expression time courses as shown in Figure 1B. Each single-cell time course consists of the mean eGFP fluorescence intensity over the square occupied by a cell during the measurement period resulting in the protein expression kinetic for the respective cell. As expected, the single-cell time courses show a large cell-to-cell variability regarding the eGFP expression (15, 16, 18). Figure 1C shows a magnification of the single-cell time courses at the beginning of the measurement illustrating the variation in the onset time points of protein expression.

### Determination of the expression onset times and expression rate

We use a kinetic biochemical rate model to describe mRNA expression to fit the individual single-cell time courses. The approach yields the individual kinetic rates and the time point when protein expression begins. As shown in Figure 2A our model includes translation of mRNA to protein and a subsequent chemical transition called maturation, which renders the GFP protein fluorescent (19, 20). In addition, the model allows for degradation of both mRNA as well as protein with time. This translation-maturation model is formulated by a set of ordinary differential equations (ODEs). The initial conditions are given by the number of mRNA molecules m_0_, which are simultaneously delivered at a defined time t_0_. The time t_0_ represents the time when translation begins and we refer to it as the onset time. The model assumes that all mRNA molecules are delivered at the same time, which clearly is a simplification. However, the time interval during which mRNA is released into the cytosol is short compared to the time scale of expression and, therefore, can be treated as a single event in time. Defining the time point of incubation as zero, the expression onset time is the period from incubation until beginning of translation, which includes all intermediated processes such as cellular uptake, release into the cytosol and unpacking of the complexes. The analytical solution to the ODEs is given in the Materials and Methods section and were used to fit the measured single-cell fluorescence time courses using least-square fitting (21). To prevent overfitting the data by using too many free parameters of the translation-maturation model we fixed the maturation rate k_M_ and the protein degradation rate β to the mean population values. These mean values were determined in an independent translation block experiment using cycloheximide performed within a previous study (22). The fitting results of the model with the fixed parameters are in good agreement with the experimental data.

**Figure 2:**
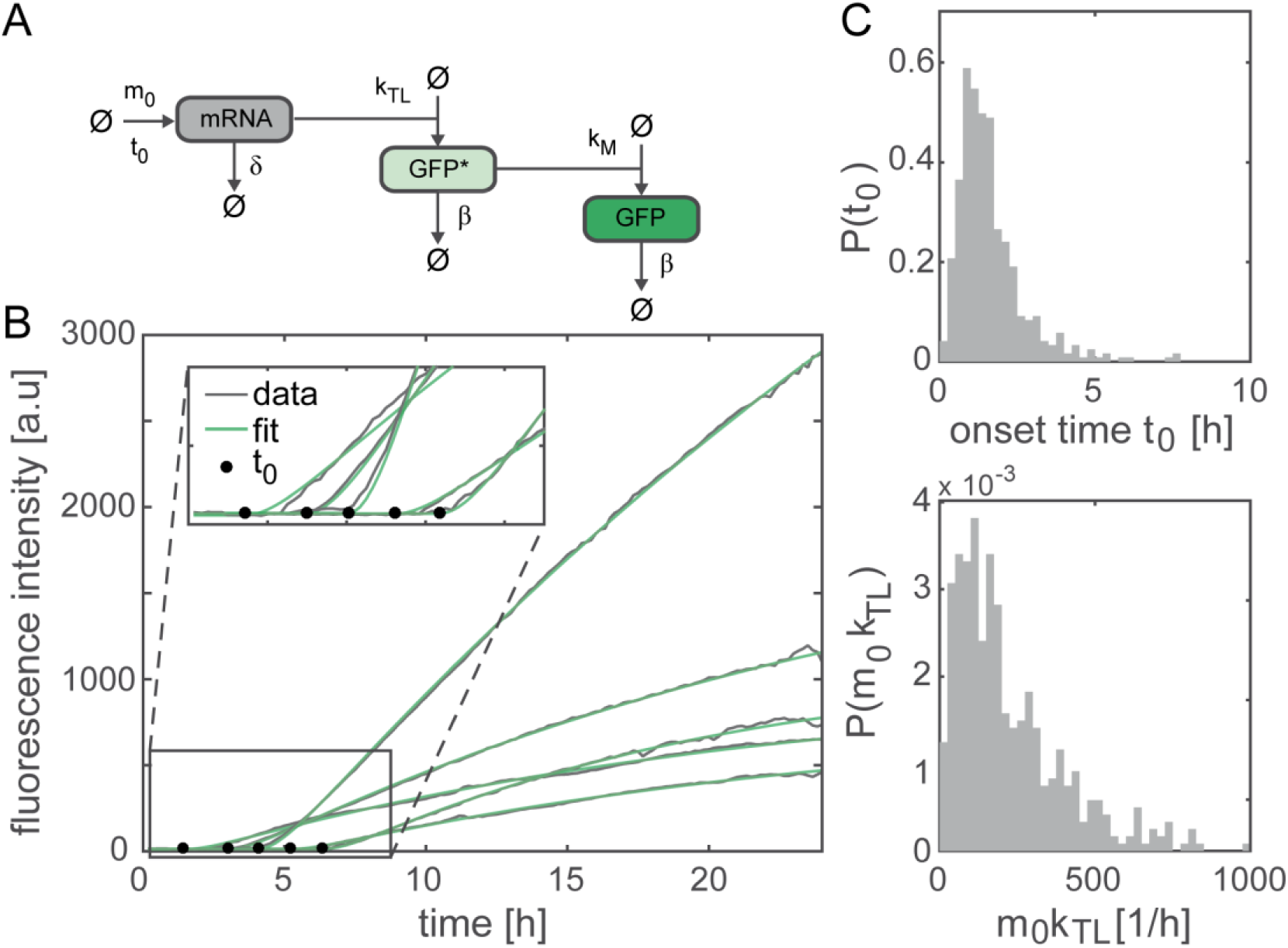
Parameter estimation of single-cell expression time courses using the translation-maturation model. (A)The translation-maturation model is sketched with the parameters plotted next to the respective reactions. (B) Representative fits (green) of the translation-maturation model to the eGFP expression time courses (gray) with fixed parameters k_M_ and β are shown in green. The insert illustrates in detail the expression onset variation and the good agreement of the fitting parameter t_0_ (black dots) with the experimental data. (C) Histograms showing the population distribution for the expression onset t_0_ and the expression rate m_0_k_TL_

Figure 2B shows representative single-cell time courses with the respective fits. The magnification of the representative data illustrates that the expression onset times (black dots in Figure2B) are reliably identified by the model. In the supplement we provide further details about the accuracy of the parameter estimation. In particular, we compare our model to alternative approaches that were explored but that we found less suitable (see Supplementary Figure 5). We analyzed the data by an automated routine, which fitted each time course by the translation-maturation model. Since maturation rate and protein degradation were fixed, the routine yields the expression onset t_0_, the expression rate m_0_k_TL_, and the mRNA degradation rate δ. The distribution of single-cell parameters of onset times and expression rates are shown in Figure 2C. The onset time show a distribution that is slightly skewed and roughly described by a gamma function (see Supplementary Figure 3A). Interestingly, the earliest observed onsets are as early as 10 min after incubation, while the later onset times extend too many hours. The average width of the onset time distribution measured as full width half maximum (FWHM) of the estimated distribution is about 2-3 hours. The expression rate is described by a log-normal distribution. The mRNA degradation rates δ also exhibit a distribution, an observation that will not be further discussed since it is outside the scope of this report (see Supplementary Figure 2).

### EGFP expression kinetics systematically change with increasing serum fraction

Next, we study the influence of nonspecific protein adsorption on the nanocarrier’s surface on the onset times and expression levels. Specifically, we increased fetal bovine serum (FBS) fractions within the medium during mRNA lipoplex incubation. To quantify if the transfection ability is systematically dependent on increasing FBS fraction we performed our assay by transfecting the cells using Lipofectamine2000 self-assembled complexes in cell culture medium with six different FBS fractions ranging from 0% to 10 % (v/v) FBS.

Combined phase contrast and fluorescence images that were taken 24 h after the cells’ transfection (see Figure 3A) show clear differences regarding the intensity of the fluorescent cells as well as the number of eGFP positive cells. The contrast settings of the combined phase contrast and eGFP fluorescence images are kept constant enabling a direct comparison of intensities. Furthermore, we used these endpoint images to determine the fraction of eGFP positive cells as measure for the transfection efficiency. The transfection efficiency decreases from 92% ± 3% for cells transfected without serum to 31% ± 6% for cells transfected with 10% FBS (data shown for cells treated with 0%, 2%, and 10% FBS in Figure 3A). Similar behavior of decreasing transfection efficiency of lipid-based vectors in the presence of serum was observed previously (23, 24). The corresponding fluorescence time courses in Figure 3B show the single-cell expression time courses together with their respective mean time course. We observed a large difference in the translation kinetics for cells transfected without serum compared to cells transfected with serum, which is in agreement with the changing transfection efficiency. The influence of serum is also reflected in the decreasing number of analyzed cells per condition, which is expected due to the decreasing transfection efficiency assuming that we have approximately the same total number of cells per condition.

**Figure 3:**
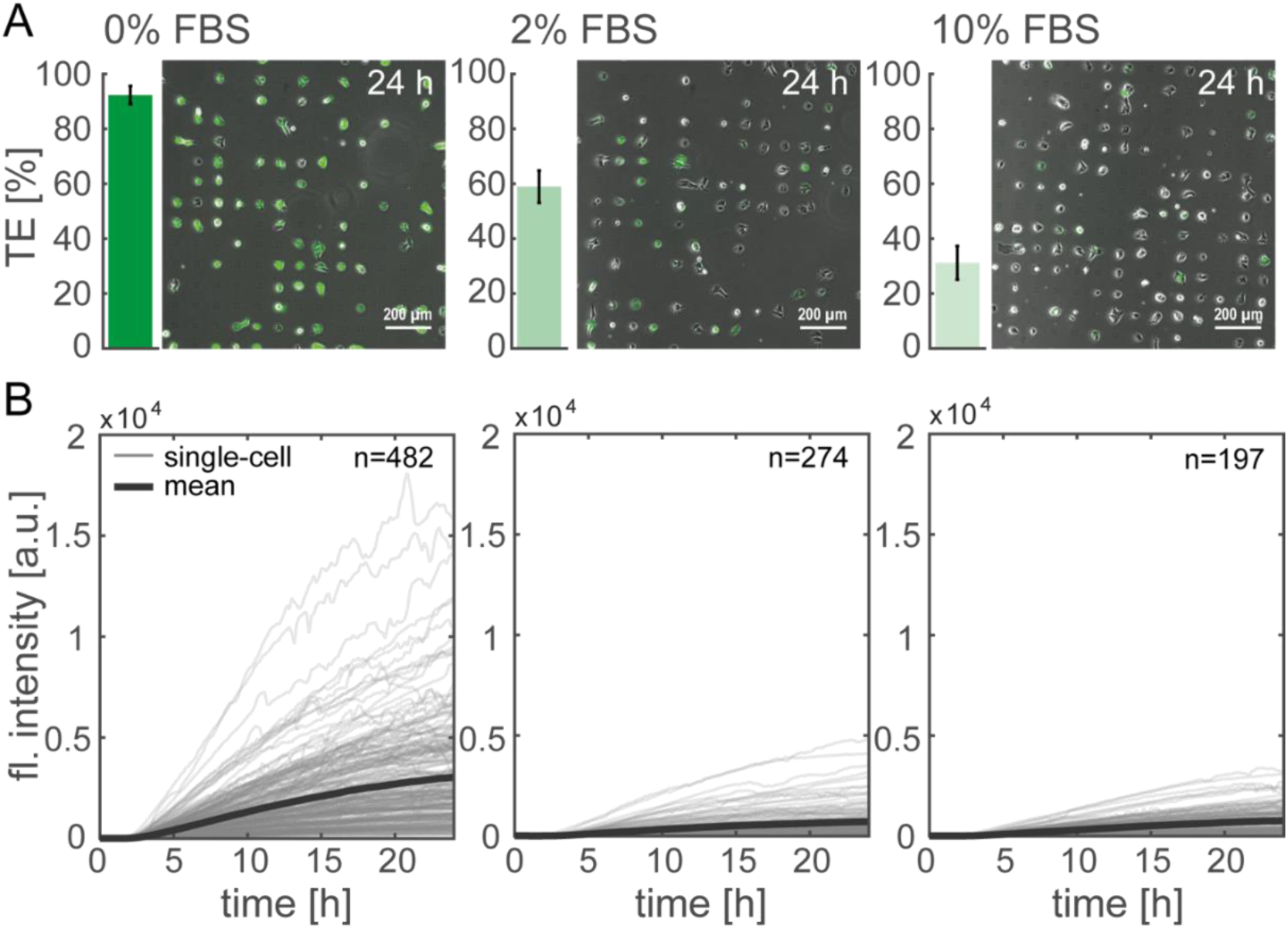
Transfection efficiency and single-cell time courses show serum concentration dependency on protein expression dynamics. (A) The transfection efficiency, defined as fraction of eGFP positive cells, decreases with increasing FBS fraction (data shown as mean ± standard deviation). The same trend is visible in the overlays showing the phase contrast images and the eGFP fluorescence signal 24 hours after mRNA transfection. (B) The single-cell time courses show the eGFP expression kinetics (gray time courses) and the corresponding mean expression (thick black time course). A clear decrease of expression level is observed between cells treated without FBS (0%) and cells treated with FBS (2% respectively 10%).

### Delivery time and expression rate distributions varies dependent on serum content

The fluorescence time courses obtained from cells transfected with varied FBS fraction are analysed by fitting the translation-maturation model to obtain values for expression onset t_0_ and expression rate m_0_k_TL_. We observed that the onset time distributions shift to later times under the presence of FBS (see Figure 4A). The mean onset time of 1.6 h for 0% FBS increases to 2.9 h for 10% FBS co-occurring with an increase of the distribution widths revealing a delayed mRNA release into the cytosol with an increased cell-to-cell variability under the presence of serum proteins (see Supplementary Figure 3A). The expression rate decreases by a factor of four between cells treated with 0% FBS and cells treated with serum, although we did not observe significant differences for the five datasets of cells treated with FBS (see Figure 4A). We expect that the decrease of the expression rate mainly reflects a decreased number of transfected mRNA molecules m_0_ and that the translation rate k_TL_ is not affected.

**Figure 4:**
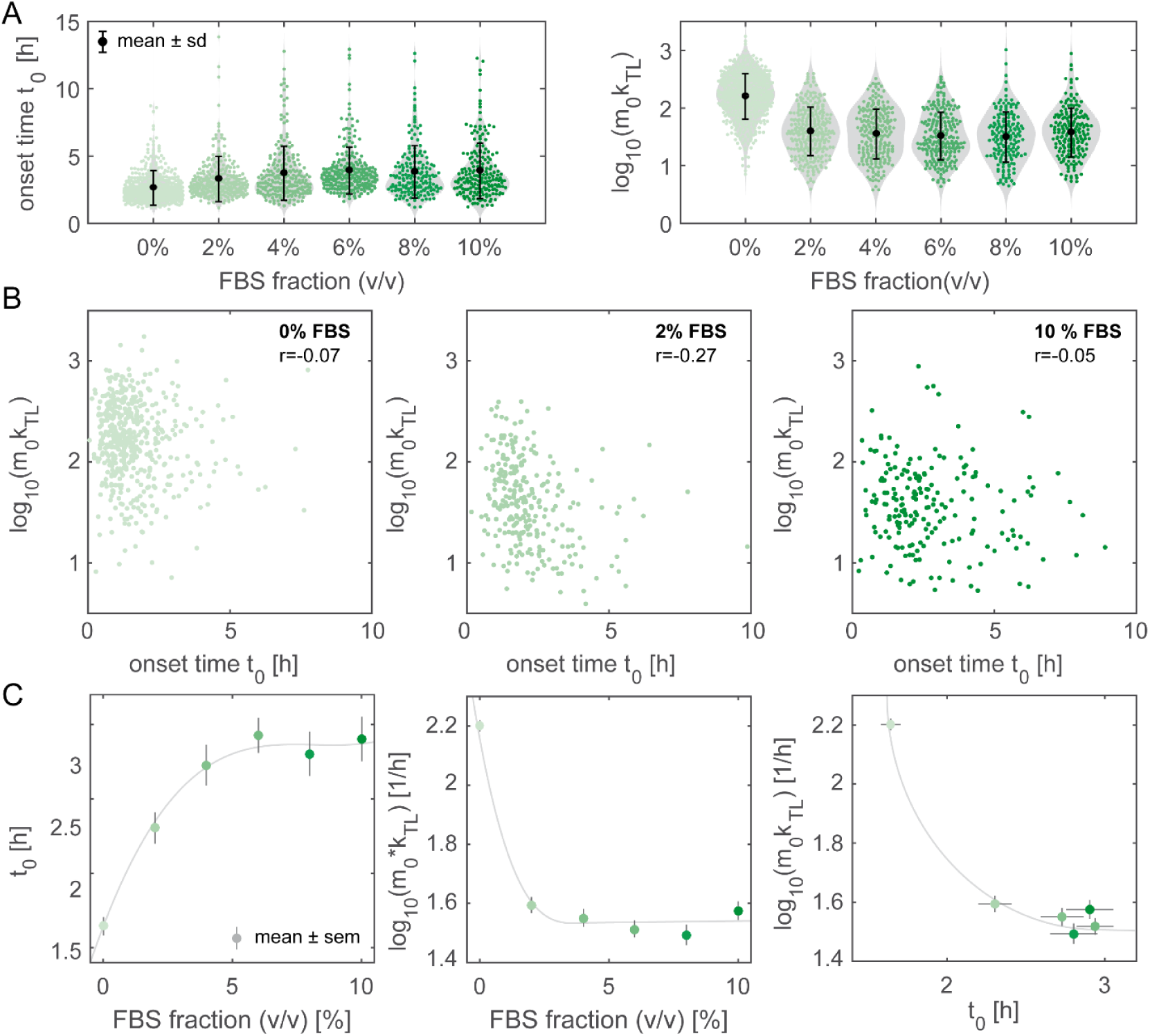
Single-cell data of onset time t_0_and expression rate (m_0_k_TL_) distributions as a function of increasing FBS fraction of the transfection medium. (A) Higher FBS contents yield in an onset time distribution shift to later times and a reduced transfection efficiency estimated using m_0_k_TL_. Each dataset of varying FBS contents shows the single-cell data as green dots with the respective mean value (full black dot) ± standard deviation. The estimated kernel densities are shown in gray to illustrate the distribution. (B) The parameters t_0_ and m_0_k_TL_are not correlated at the single-cell level. This is exemplarily shown for the datasets of 0%, 2%, and 10% FBS fraction. Each data point corresponds to a single cell and the Pearson’s correlation coefficient r is inscribed. (C) The mean onset times (± standard error of the mean) and the mean expression rate m_0_k_TL_are plotted as a function of increasing FBS content. Here, the plot expression rates versus onset times show an inverse dependence (The green data points correspond to the mean value of the six datasets with the color intensity indicating the FBS content, the gray line is to guide the eye).

To investigate whether the onset time is linked with the expression rate, we checked for correlations between these parameters. The scatter plots of the parameters, exemplarily shown for 0%, 2%, and 10% FBS in Figure 4B, are uncorrelated as indicated by low Pearson’s correlation coefficients r suggesting that there is no intrinsic mechanism connecting fast mRNA delivery with high protein expression rates within a single cell. On the other hand, both parameters show changes with increasing FBS concentrations (see Figure 4C). While the onset time increases with increasing serum, the efficiency decreases. Hence, the mean expression rates are negatively correlated with the mean onset times (last plot in Figure 4C) under the variation of the serum concentration.

### The impact of protein adsorption varies for different lipid-based nanocarriers

The Lipofectamine data show that both onset time and efficiency depend on serum concentration. However, we will demonstrate that this relation depends on the type of mRNA carrier and is not intrinsically linked. Comparing single-cell data from Lipofectamine transfection with data obtained after transfecting with LNPs made from the ionizable lipid Dlin-MC3-DMA (Materials and Methods) we observe the opposite behavior (see Figure 5). Using Lipofectamine in the presence of FBS, the distributions are shifted to longer onset times and lower expression level (see first plot of Figure 5). For cells transfected with LNPs the opposite trend is the case (see second plot of Figure 5), the onset times shortens and the expression level increases. Furthermore, we observe a more homogenous behavior for cells treated with LNPs compared to Lipofectamine treated cells, which can be seen in the less scattered LNP data.

**Figure 5:**
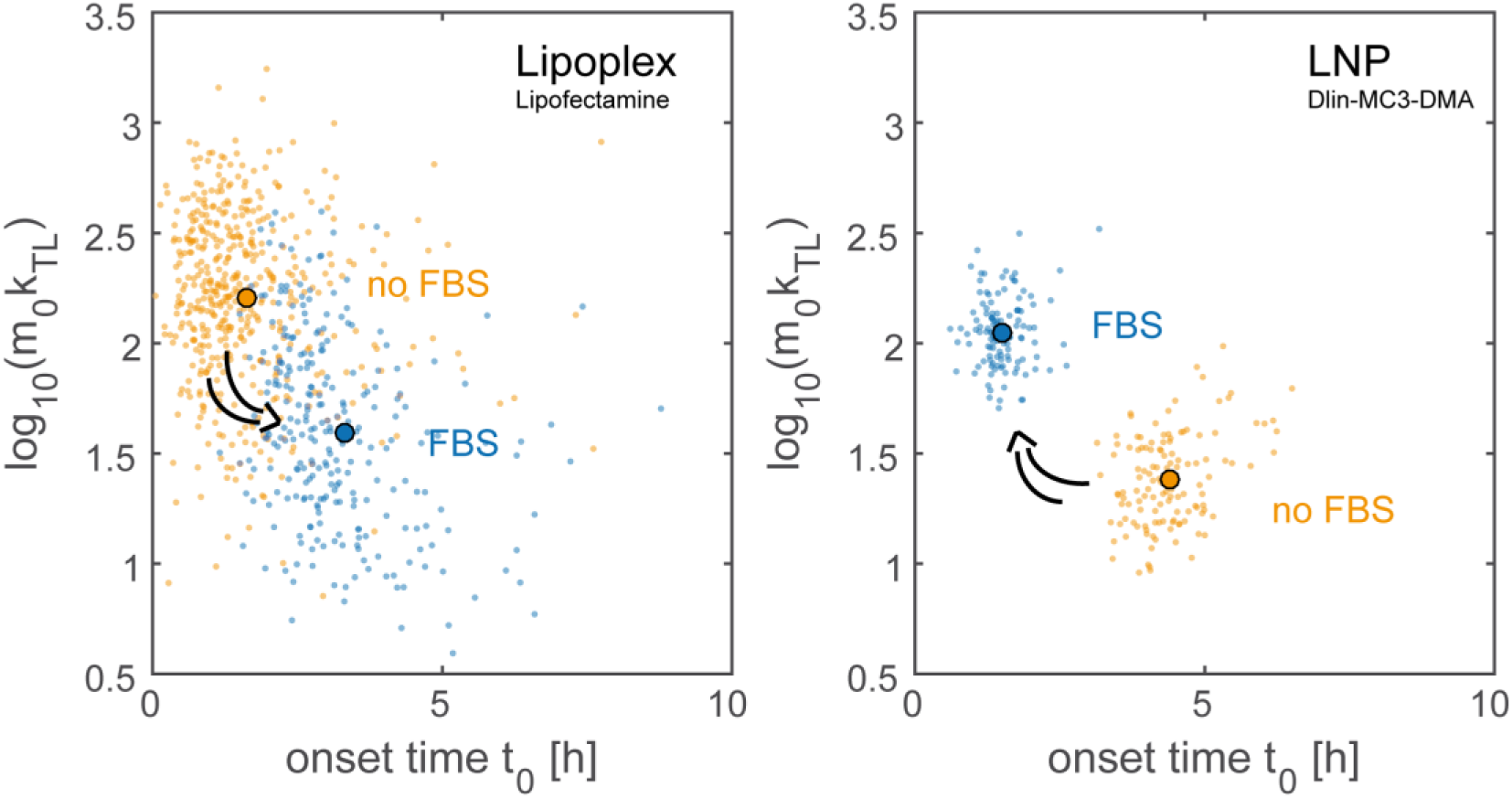
Protein adsorption on the nanocarrier’s surface has the opposite effect on lipid nanoparticles LNPs compared to lipoplexes. Scatterplots of expression onset time t_0_*vs* expression rate m_0_k_TL_ of lipoplexes (left) and LNPs (right) with (orange) and without serum (blue). The arrows indicate the opposite effect on transfection efficiency and timing induced by addition of FBS. Each data point corresponds to a single cell. The median value is indicated as full dot.

## Discussion

Using microarrays, we measured single-cell eGFP expression time courses and obtained single-cell expression rate and onset time distribution functions for mRNA transfection. The onset times display the total delivery time, which includes multiple intermediate steps consisting of mRNA nanocarrier addition to the single-cell microarray, particle adsorption on the cell surface, particle uptake, endosomal release, and the measured protein expression onset. To avoid uncontrolled long-term adsorption of complexes we used a perfusion system to apply a defined incubation period of 1 h (see Supplementary Figure 1B) during the time-lapse measurement. This enables us to cover the full dynamic range of measurable onset times including the short time scales not accessible otherwise due to need for off-microscope sample handling. In our approach, we used the expression rate as a measure for transfection efficiency. We showed that the expression rate constant m_0_k_TL_, the product of released mRNA molecules and the kinetic translation rate, is proportional to the protein level at a defined time point after transfection, which is more frequently used as the measure for transfection efficiency (25, 26). The proportionality can be explained because the expression rate corresponds to the slope of a straight line between the expression onset and a certain protein level *i.e.* 12 h after the transfection (see Supplementary Figure 6).

The delivery time distributions after mRNA transfection, are measured here for the first time at the single cell level, reveal that the delivery process occurs over a time period of only a few hours (see Figure 2C and Figure 4A). The FWHM of the distributions fitted by Gamma distribution functions increase with increasing FBS concentration (see Supplementary Figure 3A). This finding is in agreement with the literature that there is only a small window of opportunity for successful carrier release before the nucleic acid nanocarriers are trapped in late endosomes or lysosomes (12). Since we apply a pulsed incubation in our experiments, with a period restricted to only an hour, the observed onset time distribution corresponds to the delivery time distribution. Note that the onset time distribution in the case of continuing lipoplex incubation exhibits a much broader and prolonged distribution (see Supplementary Figure 7) Hence, absorption and uptake is a continuous process, while if mRNA loaded particles are trapped in endosomes or lysosomes they are subject to degradation and an intact release becomes less likely with time. This scenario explains why we do not observe late expression onsets in pulsed incubation. This mechanism, however, would suggest that cells with short delivery times show higher expression efficiency. However, we observed no correlation between delivery time and expression rate at the single-cell level (see Figure 4B). This means that there is no intrinsic mechanism linking the delivery time with expression efficiency and that the expression rates’ variance is independent of the delivery times’ variance. However, effects induced by external changes such as the level of blood serum proteins (FBS) act on both delivery time and expression rate in a systematic manner. We found that the transfection efficiency decreases, and delivery time increases for Lipofectamine mediated transfection, while the opposite occurs for LNPs. In case of Lipofectamine unspecific protein adsorption, which leads to the formation of a protein corona on the lipoplex surface, explains this effect (27, 28). The protein corona changes the structure and surface charge of lipoplexes, which influences the interaction between nanocarriers and cells for example during adsorption or endosomal uptake (29). Interestingly, the protein corona has different effects on delivery time and expression rate for LNPs, where delivery shifts to faster times and higher expression rates. It is known that different nanocarrier formulations lead to different targeting of endocytotic pathways (30) to different quantities of proteins forming the corona (31). In particular LNP formulations containing ionizable lipids and dissociable PEG lipid, the serum effect has been explained by apolipoprotein E (apoE) adsorption, (32), which has been shown to lead to increased uptake in hepatocytes via apoE receptors (33, 34). The reason why LNPs without FBS are still able to transfect might be because HuH7 cells secret apoE resulting in a lower concentration of apoE in the medium than with FBS present (35). We showed that protein adsorption on LNPs does not only play a role in faster delivery times but also influence the expression rates. However, similar to the Lipofectamine lipoplexes we did not observe a correlation of delivery time with expression rate for LNPs at the single-cell level.

Using Cy5 labeled mRNAs, we also observed that Cy5 fluorescence increased after incubation for about 1 hour. We believe that fluorescence of the Cy5 labeled mRNAs is unquenched during the time of uptake due to unpacking of the lipid-based carriers. Other studies also used multiple fluorescent signals in single cell time lapse measurement to exploit signal correlations (36, 37). Our findings are in agreement with a study by Yasar et al. showing that the kinetics of mRNA nanocarrier binding on the cell surface is the same for transfected and non-transfected cells without giving further insights on transfection efficiency (26). In future work, using fluorescent markers, which label endosomes or lysosomes, might reveal intermediate delivery steps and enable further correlation studies with endocytolytic release events.

To conclude, we presented LISCA to determine delivery times and expression rates for hundreds of cells in parallel. The approach offers insights into the correlation of nanocarrier delivery timing and efficiency. The fact that these parameters are not correlated within single cells indicates that there is no intrinsic mechanism that links the timing of delivery to the total expression rate. Additional efforts are needed to resolve the intermediate stages of nanocarrier delivery to gain a better understanding of which processes are crucial for effective delivery. Hence, the approach shows great potential as a novel method to discriminate and improve gene delivery systems with respect to delivery timing and efficiency.

## Materials and Methods

### Materials

eGFP mRNA and eGFP mRNA with Cy5 label nucleotides (996 nucleotides, 1 mg/mL in 10 mM Tris-HCl and pH 7.5) were purchased from TriLink Biotechnologies, USA. Sticky-Slide VI^0.4^® 6 channel slides and coverslips for sticky slides (uncoated) were purchased from ibidi GmbH, Germany. 100mM Sodium Pyruvate, 1M HEPES buffer, fetal bovine serum (FBS), Roswell Park Memorial Institute (RPMI) medium, Leibovitz’s L-15 medium, and the transfection reagent Lipofectamine™ 2000 were purchased from Thermo Fischer Scientific, Germany. The polymer PLL(20 kDa)-g[3.5]-PEG(2 kDa) (SuSoS Ag, Switzerland) and fibronectin (Yo Proteins, Sweden) were used for array fabrication. Sterile PBS was prepared in-house. The perfusion system for fluid handling was made of PTFE microtubing with an inner diameter of 0.3 mm (Fisher Scientific, Germany), needlefree swabable vales, female luer lugs (both MEDNET, Germany), and in-house made luer teflon plugs.

### Single-cell array fabrication

Single-cell microarrays were produced by microscale plasma-induced protein patterning (µPIPP) like already described elsewhere (38, 39). The array consists of a micropattern with cell adhesive squares, which are coated with fibronectin. The interspace between the squares is passivated with PLL-g-PEG. By cell adhesion to the squares the cells align on the micropattern enabling high-throughput read out of each cellular fluorescence signal. The whole array has six channels with the described micropattern on the bottom of each channel.

### Cell culture

The human liver carcinoma cell line HuH7 (I.A.Z. Dr. Toni Lindl GmbH, Germany) was used for the transfection experiments. The HuH7 were grown in modified RPMI containing GlutaMax supplemented with 5 mM HEPES, 1 mM sodium pyruvate, and 10% FBS. The cells were maintained in a humidified atmosphere in an incubator at 37°C and 5% CO_2_. For all transfection studies, the cells were seeded at a cell density of 10,000 cells per channel in the six-channel slide of the single-cell array 4 h prior the time-lapse measurement.

### Time-resolved fluorescence microscopy

For scanning time-lapse imaging, we used a motorized inverted microscope (Eclipse Ti-E; Nikon) with an objective lens (CFI PlanFluor DL-10x, Phase 1, N.A. 0.30; Nikon). To enable a stable temperature of 37 °C during the experiments, the single-cell array connected to the perfusion system was put in a heating box (Okolab). Fluorescence image stacks with a time resolution of 10 min were acquired using a cooled CMOS camera (pco.edge 4.2; pco), a LED light source (SOLA-SE II, lumencor), and a suitable filter cube for eGFP (BP450–490, FT510, LP 510–565; CHROMA Technology Corp.). The scanning macro defining important parameters like the exposure time or the position list was controlled by NIS-Elements Advanced Research software (Nikon).

### Transfection assay

The lipoplex solution was made by mixing Lipofectamine™ 2000 with mRNA like previously described (18). Equal fractions of this lipoplex solution were diluted to the same final mRNA concentration of 0.5 ng/µl with L15 medium containing different FBS fractions and incubated for further 5 min prior transfection.

The LNPs were fabricated by microfluidic mixing using an already published protocol (40). The LNPs contain the ionizable cationic lipid O-(Z,Z,Z,Z-heptatriaconta-6,9,26,29-tetraem-19-yl)-4- (N,N-dimethylamino)butanoate called (DLin-MC3-DMA) synthesized at AstraZeneca, Sweden. 1,2-distearoyl-sn-glycero-3-phosphocholine (DSPC) was obtained from Avanti Polar Lipids, 1,2-dimyristoyl-sn-glycero-3-phosphoethanolamine-N-[methoxy(polyethyleneglycol)-2000] (DMPE-PEG2000) was obtained from NOF Corporation, and Cholesterol (Chol) was obtained from Sigma–Aldrich. The same mRNA was used for encapsulation in the LNPs as for the lipoplexes. The formulation of the LNPs is as follows: DLin-MC3-DMA:DSPC:Chol:DMPE-PEG2000 in the ratio 50:10:38.5:1.5.

We performed the transfection during the time-lapse measurement using a tailored tubing system that was connected to the channel slide 4 h after cell seeding. First, all channels were rinsed with 37°C warm PBS. Afterwards, each nanocarrier solution was added to one channel of the single-cell array to measure the protein expression dynamics dependent on FBS fraction. After 1 h of incubation with the transfection complexes all channels were washed with L15 medium supplemented with 10% FBS, which remains in the channels for the remaining measurement. After the addition of the mRNA complexes we measured the cells for at least 15 h with a 10 min time resolution to observe the translation kinetic for each successfully transfected cell.

### Image processing and analysis

Raw images obtained after the experiments were pre-processed in ImageJ. Background correction based on (41), identification of cells on adhesion sites and readout of the total fluorescence intensity per cell was carried out using in-house written ImageJ plug-ins.

### Data fitting

We determined the delivery onset times from the fluorescence intensity data using a non-linear least-squares fit of the translation-maturation model. We used the three-stage reaction model called translation-maturation model. The model considers a translation step to produce eGFP from mRNA with rate k_TL_ while both mRNA and the protein have decay rates δ and β respectively. The translation step is followed by eGFP maturation, including folding, at rate k_M_ before becoming fluorescent. In the three-stage model we also assume that unmaturated and maturated eGFP decay with the same decay constant as the same proteases are reported for both eGFP states (42).

The differential equations for the model have an analytic solution for the amount of protein *G*(*t*) at time *t*, and this solution was fitted to the data. The analytical solution of *G*(*t*) in the three-stage model is

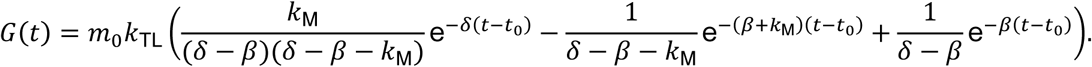

Here, *m*_0_ is the amount of mRNA taken up by the cell, *k*_TL_ is the translation rate, *k*_*M*_ is the protein maturation rate, *δ* and *β* are the degradation rates of the mRNA and of the protein (including pre-mature eGFP), respectively, and *t*_0_ is the delivery onset time relative to the exposure of cells to the transfection agent.

Due to parameter identifiability, we treated the product *m*_0_*k*_TL_ as one parameter. Moreover, since *G*(*t*) yields unphysical results for *t* < *t*_0_, we used the more robust function *Ĝ*(*t*) for fitting, which contains an additive time-independent fluorescence background parameter *z*:

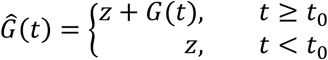

We conducted the non-linear least-squares fits using Python with Numpy and Scipy. The analysis scripts are published together with the data (see Data Availability).

For fitting the fluorescence time courses, we first estimated the starting values for the least-squares fit. The starting value for the onset time *t*_0_ is estimated by determining the time for which the fluorescence intensity has reached 10% of the total intensity range of the time series. Empirically, the fits converge best when a starting value for *t*_0_ is chosen that is later than the expected fit result. The starting value for *z* is set to the median of the first ten data points in the time course. Since the measurement started before transfection, the first data points do not contain protein fluorescence but only background. The starting values for the expression rate and the mRNA degradation rate were set to *m*_0_*k*_TL_ =1000 and *δ* =0.02, respectively. The maturation rate and the protein degradation rate were kept fixed to *k*_*M*_ =1.206707417 and *β*=0.004110525, respectively, during fitting.

This automated procedure allows us to successfully fit several hundred-time courses at once without user interaction and obtain estimations for onset times and expression rates for different transfection conditions.

## Data availability

The data and the code for conducting the analysis are available at https://doi.org/10.5281/zenodo.2626006.

## Acknowledgments

We thank A. Dabkowska and L. Lindfors (AstraZenca, Göteborg) for providing the LNPs used in this study and L. Lindfors for helpful discussion on the manuscript.

## Authors’ contributions

AR and JR conceived and designed the experiments. AR mainly performed the experiments with help of NM and RK. DW, AR, and HS analyzed the data. AR, JR, DW, and HS wrote the manuscript. All authors participated in the final editing of the manuscript.

## Additional information

### Supplementary information

Supplementary information accompanies the paper on the journal’s website.

