## Supplementary information for "Correlation of mRNA delivery timing and protein expression in lipid-based transfection"

### TIME-LAPSE MOVIE OF EGFP EXPRESSING CELLS

An example time-lapse movie is deposited under https://doi.org/10.5281/zenodo.2626006. It shows the eGFP signal of HuH7 cells on a square micropattern transfected with Lipofectamine2000 lipoplexes containing mRNA encoding for eGFP. The time stamp for each frame indicates the real time in hours. During the one hour of lipoplex addition, incubation is blended. Stacks like this were used for single-cell fluorescence time courses readout and are the basis for the data analysis of the publication.

### LIPOPLEX LABELING

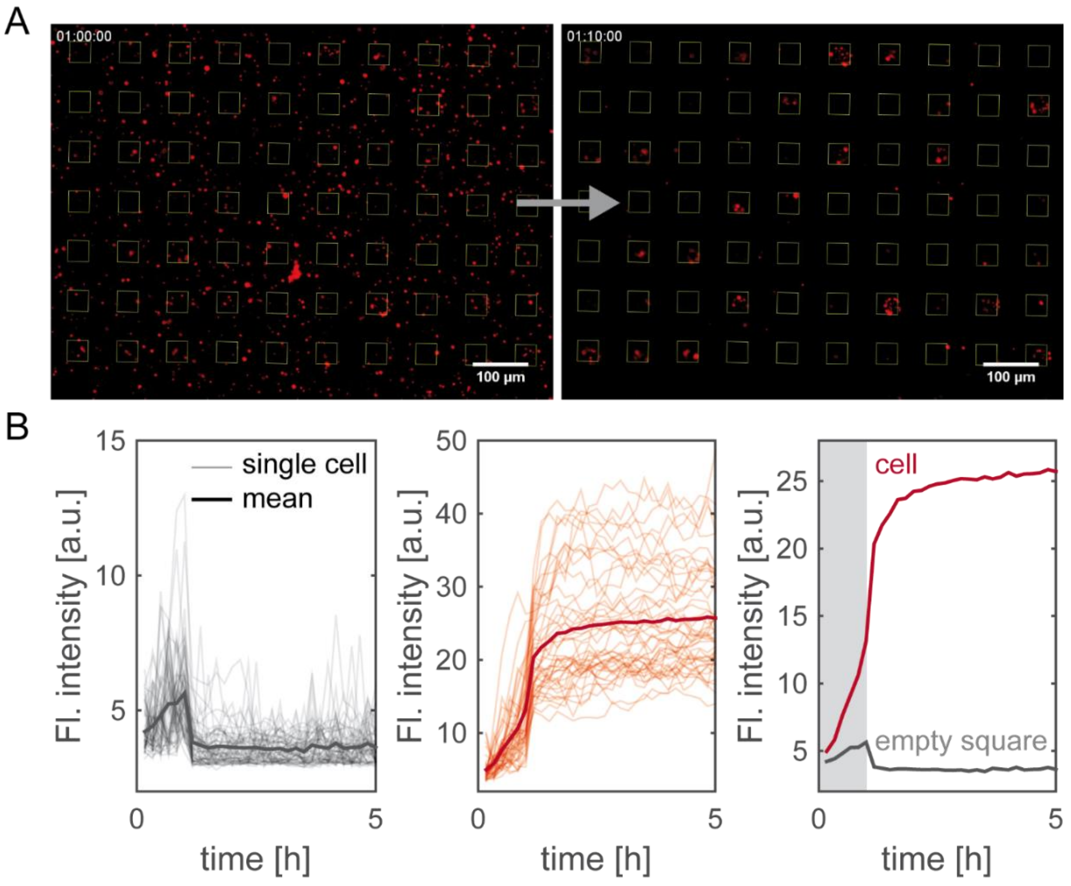


Supplementary Figure 1: Lipofectamine2000 lipoplex labeling with encapsulated Cy5 labeled mRNA. (A) The lipoplexes are randomly distributed over the micropatterened area (fibronectin squares marked in yellow) during the complex incubation (left image). After incubation the microarray is washed to flush out unbound lipoplexes. Most of the remaining lipoplexes are adsorbed on cells and are located on one of the squares (right image). (B) The Cy5 signals show clear differences between empty squares (left plot) and cell occupied squares (middle plot). The mean time course of the Cy5 signal illustrate the complex adsorption on the cell surface. The gray bar indicates the 1 h period of lipoplex incubation.

The explained single-cell translation assay of the main text can be extended with an additional fluorescence signal. As a proof of concept, we visualize the Lipofectamine lipoplexes by encapsulating Cy5 labeled mRNA (90% unlabeled mRNA and 10% Cy5 labeled mRNA). During the time-lapse measurement, we recorded the Cy5 signal as well as the eGFP expression kinetics. Supplementary Figure 1A illustrate the randomly distributed lipoplexes during the complex incubation (left image). After fluid exchange, the majority of unbound lipoplexes are removed and the remaining lipoplexes are preferentially adsorbed on cell surfaces. The positions of the fibronectin squares are illustrated by the yellow outlines. The fluorescence time courses are calculated as the mean intensity per square for each time point.

The fluorescence time courses in Supplementary Figure 1B show a weak signal increase for empty squares during the complex incubation and an abrupt decrease after the washing step at t=1 h. The squares occupied by cells show different signal dynamics (see middle plot of Supplementary Figure 1B) with comparable signal for the first hour and still increasing signal after the washing step. The single-cell time courses of the Cy5 signal show the lipoplex adsorption on the cell surface. Each of these time courses show the same behavior and just distinguish in the maximal fluorescence level dependent on the number and size of the adsorbed lipoplexes. Therefore, one can conclude that the time courses show the lipoplex adsorption process and do not contain any cell intrinsic information regarding lipoplex delivery.

### POPULATION DISTRIBUTIONS OF FITTING PARAMETERS

The four free fitting parameters of the translation-maturation model (Figure 2A) are the fluorescence offset z, mRNA degradation rate δ, protein expression onset $t_0$, and expression rate $m_0k_{TL}$. We estimated the population distribution of each parameter by fitting the model to each single-cell time courses.

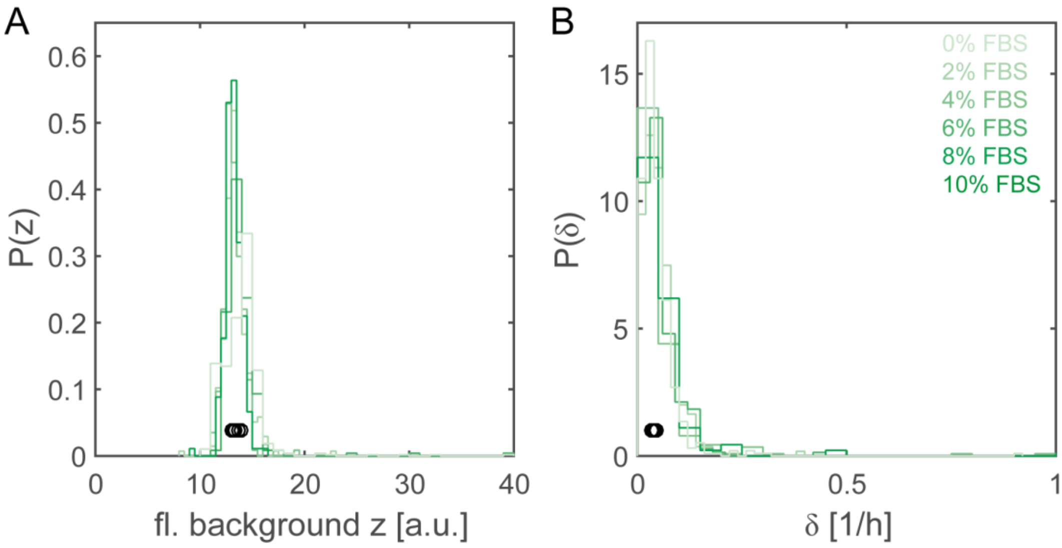


Supplementary Figure 2: The fluorescence background z and the mRNA degradation rate δ are not affected by FBS variation. The histogram outlines for each data set are shown for the different FBS fractions. The black circles show the median values of each dataset.

The distributions of the background fluorescence z (see Supplementary Figure 2A), as well as the mRNA degradation rate δ (see Supplementary Figure 2B) appear to be independent on varying FBS concentrations. The histograms for both parameters show only statistical fluctuations for the different FBS fractions with total mean values for z=13.5±0.4 a.u. and $\delta=0.052\pm0.006\ h^{-1}$ corresponding to a half-life of 13.4 h for the transfected mRNA.

The distributions of the onset time are fitted by Gamma functions (see Supplementary Figure 3A). The Gamma distributions shift to later times and the full width half maximum (FWHM) of the distributions increase from 1.8 h to 3.4 h with increasing FBS concentrations (see right plot of Supplementary Figure 3). The expression rate distributions are estimated as log-normal distributions (see Supplementary Figure 3B). The distributions for cells treated with FBS during transfection decrease to lower rates and narrower distributions. The distribution for 0% FBS is much broader with a FWHM, which is almost a factor five higher than the FWHM for cells treated with FBS.

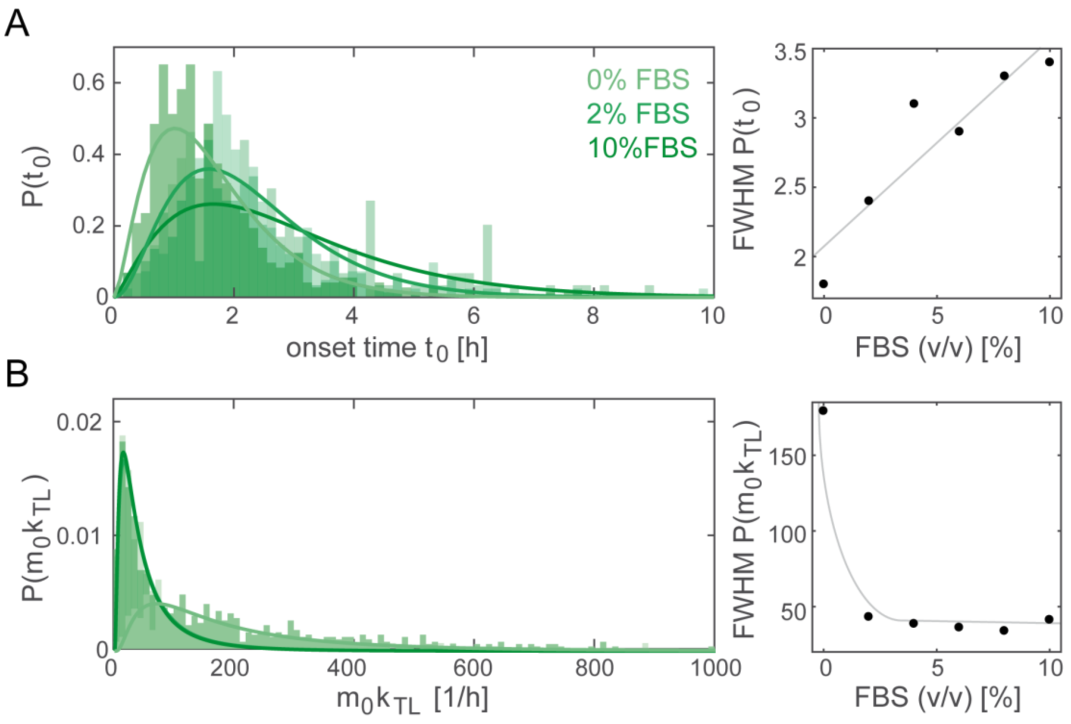


Supplementary Figure 3: The population distribution for the onset time as well as the expression rate changes under the presence of FBS. (A) The onset time distributions of all FBS datasets are estimated as Gamma distributions. (B) The distributions of the expression rate $m_0k_{TL}$ are estimated as log-normal distributions. The histograms are plotted with the respective distribution estimation. The FWHM of each distribution is plotted against the FBS fraction.

### Hierarchical Cluster Analysis for Onset Time Point Determination

In this section, we give a detailed description of the clustering-based approach for onset time extraction mentioned in the main text. Since delivery onset time detection can be regarded as a change point detection problem, where the change point is the border between the clusters of time points before and after delivery onset (1), we chose hierarchical clustering for determining the onset times.

A representative time course with the corresponding clustering is shown in Supplementary Figure 4A. Before translation onset, the fluorescence intensity fluctuates around a background level z (see blue data points of Supplementary Figure 2A). Since these fluctuations are small compared to the fluorescence of an eGFP expressing cell, the fluorescence intensity assumes values from a small and therefore densely populated interval. At the time of translation onset, the fluorescence intensity starts increasing for the rest of the measurement. Due to that persisting change, the fluorescence intensity values after translation onset are distributed with much lower density. This density difference before and after translation onset is used for finding the cluster of time points before translation onset by hierarchical clustering.

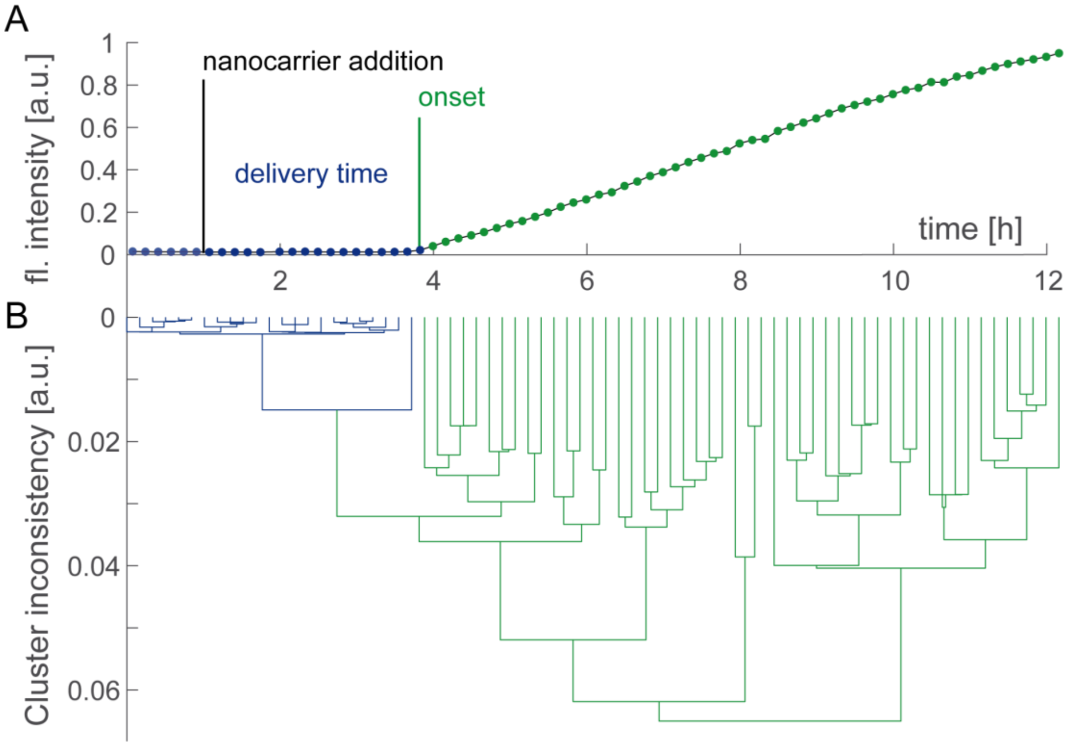


Supplementary Figure 4: A representative eGFP expression time course of a successfully transfected cell is shown. The time between the nanocarrier addition and the expression onset is defined as the delivery time. Using hierarchical clustering, the cluster of points before measureable eGFP amount is determined (blue). The last time point in the blue cluster is defined as the onset time.

The Manhattan distance is used as distance metric, which means that the distance $\mathrm{dist}(x, y)$ between two data points $x = (x_1, \ldots, x_n)$ and $y = (y_1, \ldots, y_n)$ in $n$-dimensional space is

$$\mathrm{dist}(x, y) = \sum_{i=1}^{n} |x_i - y_i| \, .$$

The distance $D(X, Y)$ between two clusters $X$ and $Y$ is calculated using the single linkage method:

$$D(X, Y) = \min_{x \in X, y \in Y} \big(\mathrm{dist}(x, y)\big)$$

A fluorescence intensity time course of a cell consists of $N$ fluorescence intensity values $f_1, \ldots, f_N$ taken at $N$ equidistant times $t_1 < t_2 < \cdots < t_N$ with $t_1 = 0$. For clustering, $n = 3$

dimensions are considered for data points at each time $t_i, i = 1, \dots, N$. These dimensions are the normalized time $\tau_i = \frac{t_i - \min_i t_i}{\gamma \cdot \left(\max_i t_i - \min_i t_i\right)}$, where $\gamma$ is a weighting factor with an empirical value of 20, the normalized fluorescence intensity value $\hat{f}_i = \frac{f_i - \min_i f_i}{\max_i f_i - \min_i f_i}$, and the time derivative of the fluorescence intensity value $d_i = \max\left(\frac{\hat{f}_i - \hat{f}_{i-1}}{\tau_i - \tau_{i-1}}, 0\right)$, where $\hat{f}_0 := \hat{f}_1$.

Using these settings, a hierarchical cluster tree is calculated, as shown in Supplementary Figure 4B, wherein the height of a link between two clusters corresponds to the distance between the clusters. The hierarchical cluster tree was calculated using Python with Scipy.

To find the initial cluster, which is the cluster of time points before eGFP onset, the initial cluster is set to the first time point and successively expanded by the next parent cluster and all its child clusters while an expansion condition is fulfilled. This is the case if: (i) the expanded initial cluster comprises at least four time points and has a height of at least 0.006; (ii) if the expanded initial cluster has a height not above 0.025 and includes time points before the latest time point of the previous initial cluster that did not belong to the previous initial cluster; or (iii) if the expansion step only adds time points before the latest time point of the previous initial cluster. All constant parameters used in evaluating the expansion condition are chosen for empirical reasons.

When the expansion condition is not fulfilled any more, the time corresponding to the latest time point within the initial cluster is identified as the expression onset time. As for the translation-maturation model, the onset time is determined relative to the nanocarrier addition to measure the delivery time.

### COMPARISON OF EXPRESSION ONSET DETERMINATION

Beside the clustering approach and the used translation-maturation model, we tested a two-stage model for data fitting. We compared the two-stage model with the three-stage reaction model, which is the translation-maturation model explained in the main text. The two-stage model only considers a single translation step to produce eGFP from mRNA with rate $k_{TL}$ while both mRNA and the protein have decay rates $\delta$ and $\beta$ respectively (2-4). The three-stage model takes into account that eGFP undergoes maturation (complete folding at rate $k_M$) before becoming fluorescent. In the three-stage model we also assume that unmaturated and maturated eGFP decay with the same decay constant (5).

The differential equations for both models have analytic solutions for the amount of protein $G(t)$ at time $t$, and these solutions were used to fit the data. The analytical solution of $G(t)$ for the two-stage model is

$$G(t) = \frac{m_0 k_{\mathrm{TL}}}{\delta - \beta}\left(\mathrm{e}^{-\beta(t-t_0)} - \mathrm{e}^{-\delta(t-t_0)}\right).$$

Supplementary Figure 5A shows representative examples of single-cell fluorescence time courses and their corresponding best fits for both models and the respective onset times for all three approaches. As can be seen in Supplementary Figure 5A, the two-stage translation model (green time course) implements the delivery onset as an abrupt kink and therefore systematically overestimates the onset times, whereas the 3-stage maturation model reveals the smooth onset behavior exhibited by the data. This difference is also reflected in the distributions of onset times in Supplementary Figure 5B. The values obtained using the model-free clustering approach typically lie between the results of the two model-based approaches and are mostly closer to the respective 3-stage onset time than to the 2-stage onset time (see Supplementary Figure 5). The hierarchical clustering approach has the disadvantage that the expression rate $m_0k_{TL}$ cannot be determined and can only be substituted by the fluorescence intensity level at a distinct time point (see Supplementary Figure 6).

We concluded that protein maturation must not be neglected in models for a reliable estimation of expression onset times, and that the 3-stage translation-maturation model provides an accurate description of the expression time course including the onset dynamics.

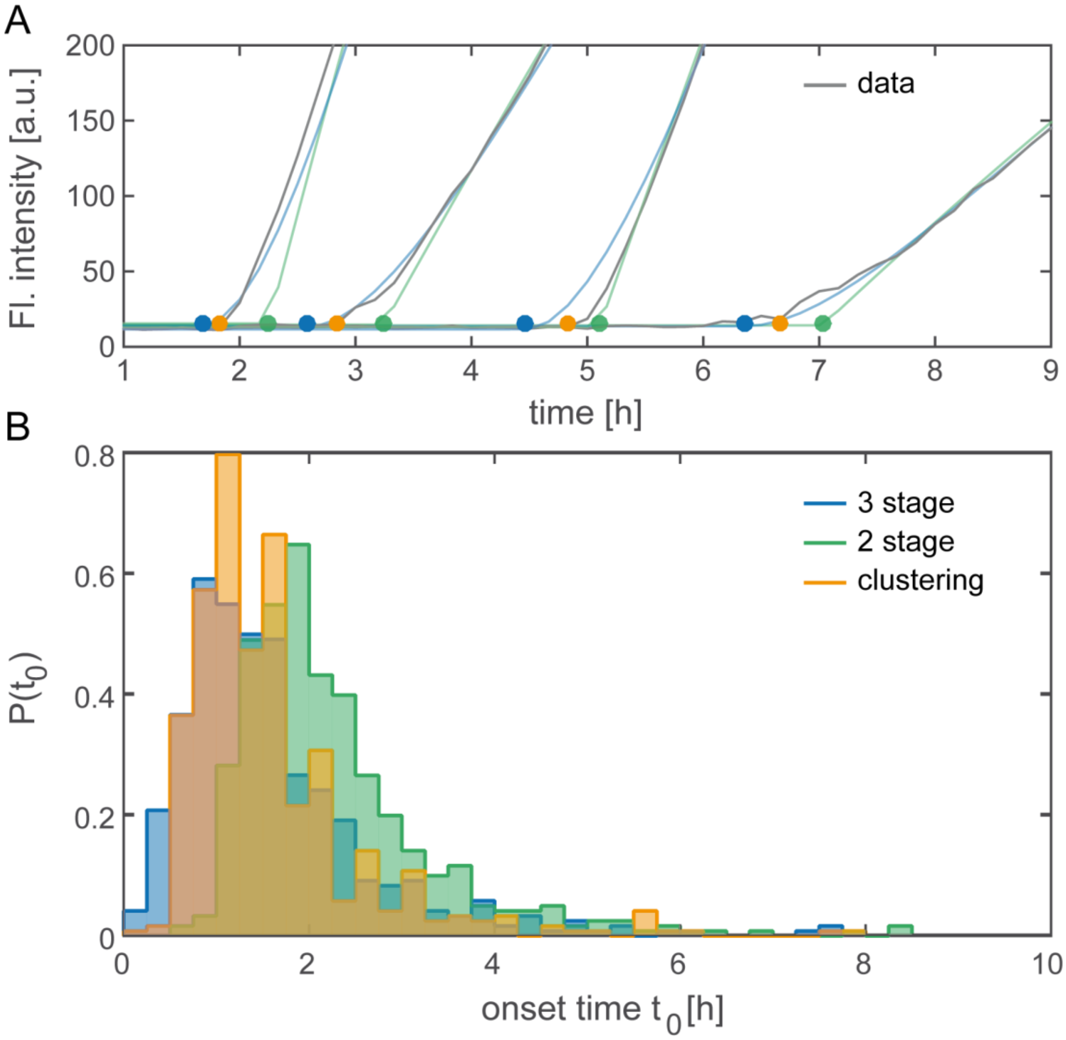


Supplementary Figure 5: Comparison of the three different methods for determination of the onset time $t_0$. (A) Representative time courses (gray) with the respective fits for the two-stage (green) and three-stage (blue) model. The onset time points determined by the models and the clustering approach (orange) are illustrated as dots. (B) The distribution of the two-stage onset times (green) is shifted to later time points compared to the other two approaches. The clustering (orange) onset time distribution is similar to the three-stage (blue) distribution.

### DETERMINATION OF TRANSFECTION EFFICIENCY

In the main text the expression rate $m_0k_{TL}$, the product of released mRNAs $m_0$ with the translation rate $k_{TL}$, is used to define the transfection efficiency for each transfected cell. Using the model independent clustering approach to determine the onset time distribution it is not possible to determine $m_0k_{TL}$. However, the fluorescence intensity 12 h after nanocarrier addition can be easily determined as a good approximation for $m_0k_{TL}$. The scatterplot of these two parameters in Supplementary Figure 6B shows a strong correlation (Pearson's correlation coefficient of 0.94) proofing that the fluorescence intensity value 12 h after transfection is a good approximation for the expression efficiency. This strong correlation can be explained by the fact that expression rate can be understood as the slope of a straight between the onset time and the fluorescence intensity 12 h after transfection (see Supplementary Figure 6A).

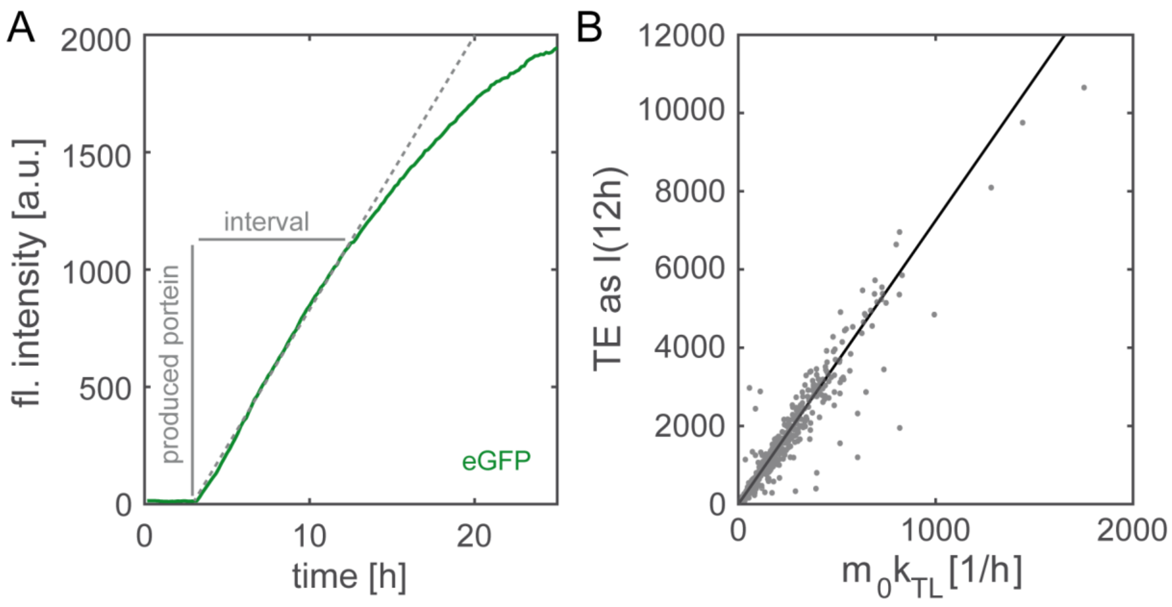


Supplementary Figure 6: (A) The translation rate $k_{TL}$ is proportional to the amount of produced protein within a defined time interval and correspond to the slope of the dashed line. (B) The transfection efficiency TE measured as the fluorescence intensity at 12 h I(12h) is strongly correlated with the expression rate at the single-cell level as visualized by the black straight.

### ONSET TIME DISTRIBUTION SHIFTS FOR LONG-TERM LIPOPLEX INCUBATION

We discussed in the main paper the use of a perfusion system to transfect cells on stage during the scanning time-lapse measurement. The perfusion system is an advantage for two reasons. Firstly, early fluorescence intensity changes after transfection can be monitored, which could not be detected if the time-lapse need to be set up after off-microscope transfection. Secondly, the period of lipoplex incubation can be controlled by flushing out unbound lipoplexes after a pulse like incubation period of 1 h. The transfection on stage leads therefore to a direct measurement of all onset times including the very early ones and eliminates the possibility of late lipoplex adsorption after the washing step. The pulse like incubation is important to assure that the lipoplexes are all adsorb within a short time frame, which enables the assumption that the protein expression onset correspond to the delivery time. This assumption cannot be made if the lipoplex solution remains on the cells for the whole measurement. For long-term Lipofectamine incubation, the onset time distributions shift to later times compared to cells incubated for only 1 h (see Supplementary Figure 7). This leads to a change of the mean value from 1.6 h for the 1 h incubation to 3.5 h for the long-term incubation.

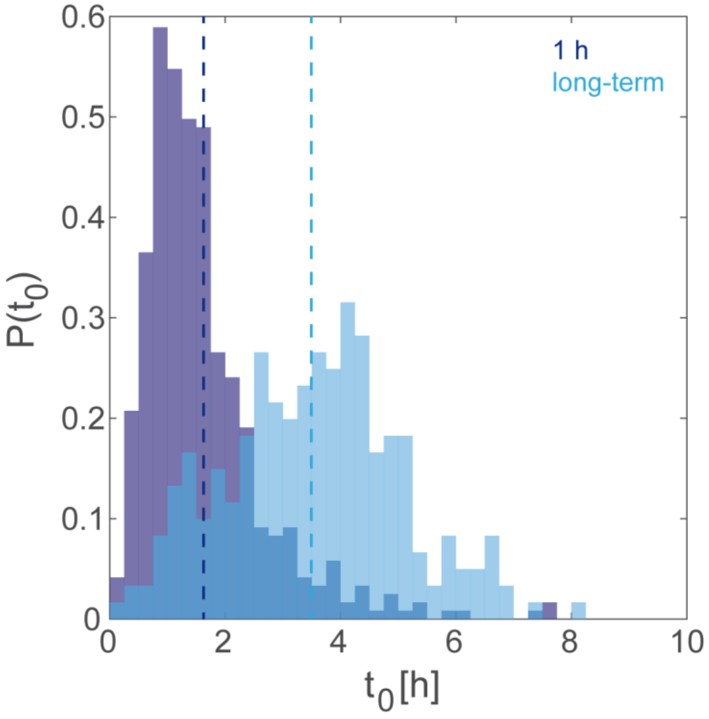


Supplementary Figure 7: The onset time distribution shift to later times for long-term lipoplex incubation compared to controlled incubation period of 1 h. The mean value for each distribution is shown as dashed line.
